# Soil microbiome predator diversity outperforms nitrogen addition in boosting plant biomass via bacterial community shifts

**DOI:** 10.64898/2025.12.02.689768

**Authors:** Alejandro Berlinches de Gea, Joshua Both, Nina Haas, Rutger A. Wilschut, Florian Wichern, Stefan Geisen

## Abstract

Nitrogen (N) is crucial for plant growth, but its overuse harms biodiversity. Increasing soil biodiversity might provide the means to reduce N inputs, but experimental evidence for this paradigm-shift is limited. Using microbiome predators (protists and nematodes) that shape microbiome composition and release N, we examined how interactions between their diversity and N addition affect *Cannabis sativa* growth. Microbiome predator addition overall boosted plant biomass by up to 53%, particularly under low N conditions, primarily by altering bacterial community composition and enriching functions related to carbon and N cycling. In turn, microbiome predator diversity had the strongest effect on biomass production, while N levels played a greater role in determining plant nutrient content. These findings suggest that increased microbiome predator diversity can reduce the plant’s dependency on added N, showing the pivotal role of soil biodiversity in enhancing plant performance and serving as tools to mitigate N inputs.

## Introduction

Inorganic nitrogen (N) fertilization was part of the green revolution and ever since provided the basis to feed the growing human population [1]. However, crops in agricultural systems take up only 60% of added N and N use efficiency declines with agricultural intensification [2, 3]. The remaining N leaches out of agricultural systems and negatively affects the environment, such as through the accumulation of nitrate in groundwater, soil acidification, and increased greenhouse gas emissions [3, 4]. Excess N also reduces biodiversity such as plant and insects [5, 6], by favoring opportunistic ruderal and pest species that thrive under increased N availability [7].

The loss of biodiversity, as explained by the biodiversity and ecosystem functioning (BEF [8]) concept, has far-reaching consequences as a reduced biodiversity (here: species richness) is commonly associated with a loss in ecosystem functions. For instance, in a global study, Hautier et al [9] showed that grassland communities with more diversity exhibited a higher multifunctionality. While mainly shown for plants, this BEF relationship also has been proven for different organismal groups across ecosystems [10–12] and is explained by the increasing complementarity of functions in the multidimensional niche space [9, 13]. However, the soil BEF (sBEF) concept so far showed relationships to range from positive to negative, depending on abiotic conditions [14].

The little attention given to sBEF compared to BEF contradicts the fact that soils host 59% of the total biodiversity on our planet [15], determining essential ecosystem functions like carbon and N cycling or plant growth [16, 17]. Among these ecosystem functions, bacterial-driven N cycling is particularly important [18–20], especially in N-limited soils [21, 22], while in agricultural soils plant’s dependency on naturally occuring N-increasing processes has been removed by fertilization. However, the role of bacteria in ecosystem functioning may be affected by microbiome predators, which are key drivers of bacterial community composition.

These microbiome predators (here: protists and nematodes) release nutrients immobilized in bacteria and fungi through predation, enhancing N availability via the microbial loop [23–25]. By selective feeding, microbiome predators shift bacterial communities [26], influencing functions like N and C cycling [23, 27]. However, most studies on species-specific prey selection focus on single model microbiome predator species [28, 29] and few have experimentally manipulated microbiome predator diversity to test sBEF relationships [30]. Thus, despite their diversity and functional importance, the effects of increasing microbiome predator diversity on plant performance, underlying mechanisms, and potential interactions with N availability remain unclear.

We explore how increasing soil microbiome predator diversity interacts with varying N addition to affect plant performance (*Cannabis sativa* L.). To do so, we inoculated pots containing a sand-potting soil mixture with four predator diversity levels (0 to 20 protist species + 0 to 4 nematode species) across four N levels (0 to 150 kg N ha□¹ yr□¹). To provide a broader picture of the sBEF relationship we correlated several plant and soil parameters such as soil microbial biomass C and N, as well as total N in the soil, plant nutrient content (here used to integrate the measurements of chlorophyll content, leaf C, leaf N, leaf S and leaf C/N ratio) and identified bacterial diversity by amplicon sequencing. Additionally, we used Structural Equation Modelling (SEM) analysis to identify potential underlying mechanisms linking microbiome predator diversity to plant performance. We hypothesize that 1) The sole addition of microbiome predators, irrespective of diversity levels, will increase plant biomass and nutrient content, especially at low N levels. 2) An increasing diversity of microbiome predators will increase plant performance, especially at low N levels. 3) Changes in plant biomass due to predator addition/diversity can be explained by shifts in plant and soil nutrient parameters as well as by the bacterial community composition.

## Materials and Methods

### 1. Experimental set-up

The greenhouse pot experiment was carried out at Wageningen University & Research, the Netherlands. It included a fertilization treatment with four levels (0, 25, 100 and 150 N kg ha^−1^ year^−1^) crossed with four microbiome predator diversity levels (0, 6, 12, and 24 microbiome predator species). Each of the 16 treatment combinations was replicated 10 times, resulting in 160 1L pots. The soil in the pots was composed of a 1:1 mixture of sieved (2 mm) river sand and low-nutrient potting soil (Lensli, Bleiswijk) previously autoclaved (121–136 ◦C, 4.5 h, 2.5 bar; Scholz, Germany). While N regimes were chosen based on agricultural practices of hemp cultivation (*C. sativa* cv. ‘Ivory’), an initial amount of 270 kg of total N ha^−1^ year^−1^ was already present in the soil before starting the experiment. This was done by measuring the N content in the soil used previous to the experiment, as described in section 8. N was added to the pots using ammonium nitrate (NH_4_NO_3_) (VWR Chemicals, Pennsylvania) eight days after inoculation with the microbiome predators (See 2.2 for details on microbiome predators inoculation). The diversity levels were created as follows: 0 species of protists and nematodes; 5 species of protists and 1 nematode species; 10 species of protists and 2 of nematodes; 20 species of protists and 4 of nematodes. Pots containing 0 species of microbiome predators were used as a control for the diversity levels and pots with 0 microbiome predators and 0 N kg ha^−1^ year^−1^ were used as control for all treatments (Fig. 1).

**Figure 1.**
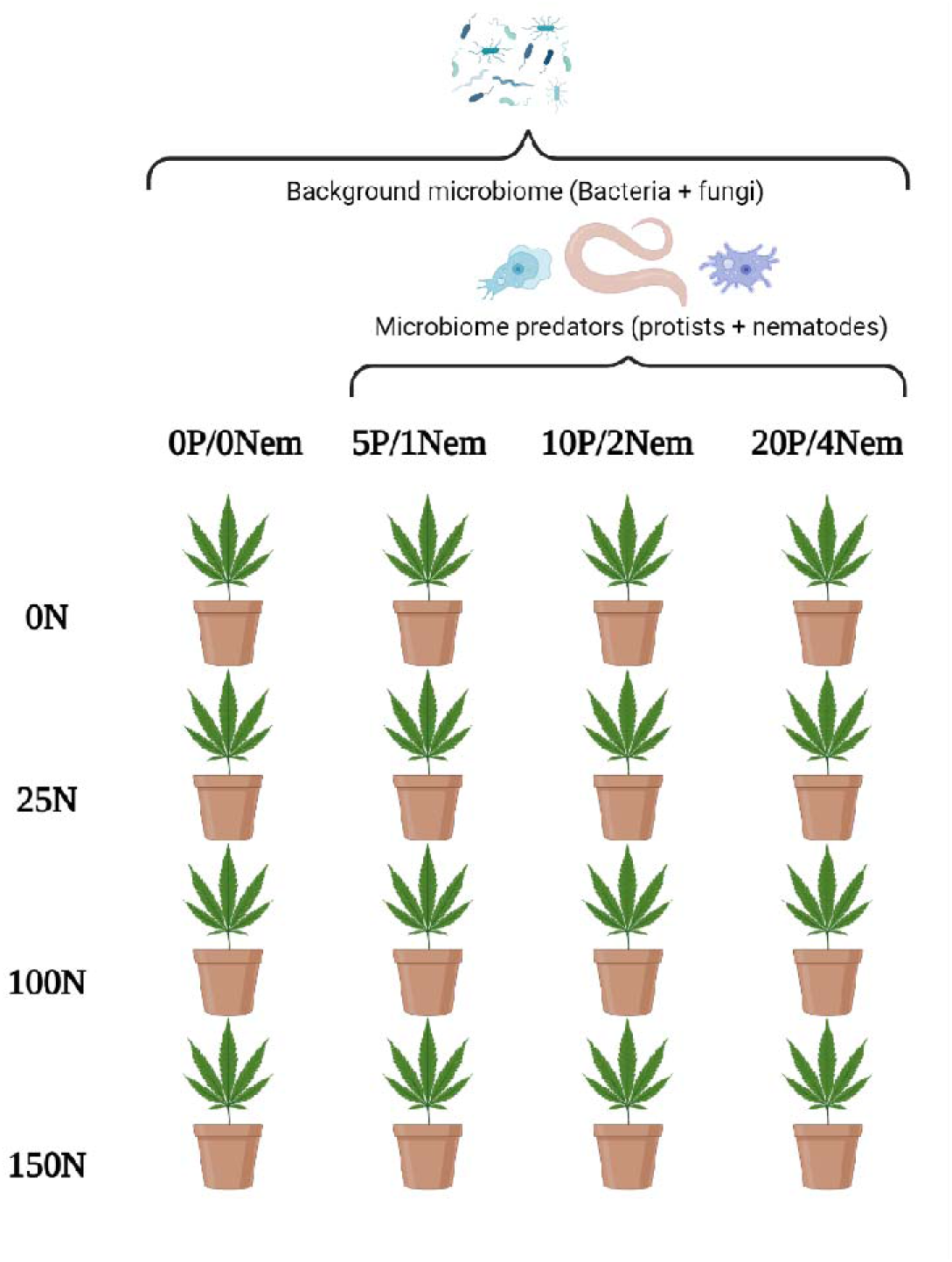
Experimental design detailing the components of each treatment. On top, the background microbiome inoculated to all the pots and the four levels of biodiversity are depicted. All treatments were replicated 10 times. The pots containing 0 protist (P) and 0 nematode (Nem) species under no N regime were used as a general control for all the treatments while pots with 0 protist, 0 nematodes, and N regimes were used as a control within treatments.

### 2. Inoculation of soil biota

#### 2.1 Microbiome treatment

Suspensions of a standardized, diverse community of bacterial and fungal cultures were added into plastic bags containing the soil used for the experiment (See section 3.1 for suspension preparation details). These bacterial and fungi suspensions were divided over the bags containing soil proportional to the bag’s weight, using an accu-jet pro pipette (Brand, Wertheim). As there were 30 bags of soil varying between 7.5 and 12.5 kg, every bag was inoculated with 15-25 ml of bacterial suspension and 25-40 ml of fungal suspension accordingly. The same soil was used to fill up the germination trays (see section 4 for germination conditions) and the pots that hosted the seedlings for the rest of the experiment.

#### 2.2 Microbiome predator treatment

Protists and nematodes (See 2.3.2 for microbiome predator isolation and cultivation) were added to the pots 48 h after the seedlings were transferred to the pots (see section 4 for germination conditions). To maintain consistent abundance (approx. 1700 protist and 7000 nematodes per pot) across the three diversity treatments, both protists and nematodes were added in a 1: 0.5: 0.25 volume ratio among the treatments (5, 10, 20 protist species; 1, 2, 4 nematode species) using a regular pipette. With this setup, protist and nematode species richness varied, but abundances remained equal among treatments. Each microbiome predator community added was randomly assembled, ensuring that each replicate of the treatment had a unique set of microbiome predator taxa, thus avoiding keystone species effects [31]. By creating one distinct microbiome predator community per replicate, we generated 40 different communities which were the same for the four N treatments. Consequently, each protist and nematode species were roughly equally distributed across all treatments, with each protist species present in 60-64 pots and 68-72 pots for nematodes. To account for the varying volumes of added suspensions, an equal amount of NMAS (the same buffer used to inoculate the microbiome predators) was added to the pots that did not receive any microbiome predators.

### 3. Preparation of soil biotic treatments

#### 3.1 Microbiome treatment

For bacteria, we used the same cultures created and used by Berlinches de Gea et al [30]. Shortly, the cultures were extracted from seven different soils located in the surroundings of Wageningen (51◦58′18″ N 5◦41′02″ E; 51◦58′44″ N 5◦42′15″ E; 51◦58′23″ N 5◦43′05″ E; 51◦58′57″ N 5◦43′26″ E; 51◦57′56″ N 5◦38′33″ E; 51◦57′16″ N 5◦36′16″ E; 51◦57′26″ N 5◦36′08″ E) according to Geisen et al [32]. Before inoculation, bacterial suspensions were adjusted to 0.1 optical density (OD600) using a Pharmacia Novaspec II spectrophotometer (Pharmacia Biotech Novaspec II, The Netherlands). Lastly, all subcultures were pooled into the same 500mL Schott Duran glass bottle (DWK Life Sciences GmbH, Germany).

Fungal cultures used in this experiment were a subset of the cultures used by Berlinches de Gea et al [30] (*Acremonium* sp., *Alternaria alternata*, *Arthrinium phaeospermum*, *Chaetomium globosum*, *Fusarium oxysporum*, *Gibberella avenacea*, *Ilyonectria macrodidyma*, *Mucor hiemalis*, *Neonectria radicícola*, *Penecillium chrysogenum*, *Plectosphaerella cucumerina*, *Tricoderma citrinoviride*). These cultures were kept in potato dextrose agar (PDA) containing 1 L demineralized water, 19 g of potato dextrose agar powder (OXOID, UK) and 7.5 g Select agar powder (Invitrogen, UK). We created a fungal suspension by scrapping the entire surface off with 1 mL NMAS using a cell scraper (Greiner Bio-One., The Netherlands). As for bacteria, all subcultures were pooled into a 1L Schott Duran glass bottle (DWK Life Sciences GmbH, Germany).

#### 3.2 Microbiome predator treatment

Protist species used were a subset of the species cultures used by Berlinches de Gea et al [30] (*Acanthamoeba castellanii*, *Acanthamoeba* sp., *Allovahlkampfia* group 1, *Allovahlkampfia* group 2, *Cercomonas lenta*, *Cercomonas* sp., *Cochliopodium minus*, *Didymium* sp., *Heterolobosea* sp. 1, *Heterolobosea* sp. 2, *Mycamoeba* sp., *Naegleria clarki*, *Naegleria* sp. 1, *Naegleria* sp. 2, *Rosculus terrestris*, *Spumella* sp., Unidentified ciliate, Unidentified cilliate 2, Unidentified cilliate 3, Unidentified flagellate, *Vahlkampfia bulbosis*, *Vannella* sp., *Vannella* sp. 2, *Vermamoeba vermiformis*). Shortly, protists were extracted from soils using a modified liquid aliquot method (LAM) as described by Geisen et al [33].

In addition, we used four species of bacterivorous nematodes (*Acrobeloides varius*, *Caenorhabditis remanei*, *Rhabditophanes* sp. and *Rhabditis* sp.) obtained from cultures available in the laboratory of Nematology, Wageningen University & Research. These cultures were grown in a Nematode Growth Medium containing 3g NaCl, 2.5g bacterial peptone, 17g Agar, 975mL water, 1mL cholesterol, 1mL 1M CaCl_2_, 1mL 1M MgSO_4_, and 25mL 1M potassium phosphate and kept at 20°C. *Escherichia coli* (OP50) was added to each plate to feed the nematodes. To reach nematode numbers, 2-3 small pieces of the NGM cultures were transferred to a new NGM every 10 days. Before the inoculation, nematode cultures were washed with NB-NMAS[34] liquid media, the same media used to maintain protist cultures.

### 4. Seed germination

*Cannabis sativa* (variety ‘Ivorý) seeds were sown into a plastic-covered tray containing autoclaved soil (121–136 □C, 4.5 h, 2.5 bar; Scholz, Germany) inoculated with bacteria and fungi (See section 2.1 for inoculation details). This soil did not contain microbiome predators. The plastic remained for 1 week to ensure the right germination conditions. After two weeks of growth, seedlings were planted in 1L pots inoculated with the same bacterial and fungal community and composed of a 1:1 mixture of sieved (2 mm) and autoclaved (121–136 □C, 4.5 h, 2.5 bar; Scholz, Germany) river sand and low-nutrient potting soil (Lensli, Bleiswijk).

### 5. Greenhouse conditions

Plants were grown for 42 days with a soil moisture content kept around 50% of the water-holding capacity and air moisture content at 70%. The plants were exposed to a day/night regime of 16/8 h of artificial photosynthetically active radiation daylight (PAR, 400-700 nm) and 18/20 □C. Plants were randomly rearranged every two weeks to avoid an effect caused by environmental heterogeneity in the greenhouse.

### 6. Harvest and processing of plant and soil samples

After 42 days of plant growth, plants and soil were harvested. Watering was stopped 4 days pre-harvest to ease the process of soil removal from roots. Stems and leaves were cut off at the soil level using plant scissors. Plant biomass was measured after drying at 70 □C for 120 h. The root system was separated from the soil by gently shaking while aiming to reduce root damage. Roots covered with soil particles were stored at 4 □C, after which they were washed. Root biomass was then measured after drying for 48 h at 70 □C. To determine soil moisture, 5-10 g of soil was collected, dried at 70 □C for 144 h and re-weighed. For pH measurements, 5-10 g of soil was suspended in a 2:5 ratio with demineralized water in a 50 mL falcon tube (Sarstedt; Numbrecht, Germany). The falcon tube was shaken for 2 h at 250 rpm (Multifuge 3 S-R Refrigerated Centrifuge; Heraeus, US), after which pH was determined directly from the liquid using a soil pH meter with a glass electrode (SI analytics, Mainz). Soil samples were stored at 4 □C for posterior nematode extraction and at −20 □C for DNA extraction (see section 7). An extra 100 g of soil per sample was stored at 4 □C to conduct analyses for microbial biomass carbon (MBC), microbial biomass nitrogen (MBN) and available NH_4_^+^ NO_3_^−^ content (see section 8).

### 7. Nematode counting, soil DNA extraction, sequencing and qPCR

Soil nematodes were extracted using Oostenbrink elutriation [35], stored in a glass 100 mL jar at 4 □C and counted in a petri dish with gridlines (50 mm x 15 mm) under an IXplore Standard microscope at 100x magnification (Zeiss, Stuttgart) to determine potential cross contaminations. In fact we found nematodes in pots that were not inoculated with microbiome predators, indicating contaminations and hindering the real potential implications of our results. Nevertheless, the significant differences found among treatment levels suggest that the initially inoculated microbiome predator communities were efficiently established and impacted plant performance throughout the experiment, aligning with the priority effect theory [36].

DNA was isolated from 1g of soil as described by [37] with the only difference of using 1.5 g of silicon carbide powder (ThermoFisher, Kandel, Germany) instead of 3ml of bead solution and iron beads. The extracted DNA samples were stored at −20 □C. DNA concentration was then quantified with a Qubit® Fluorometer device (Thermo Fisher, USA) by using a dsDNA Broad Range kit (Thermo Fisher, USA) and sent to Genome Quebec (Quebec, Canada) for 2×300bp pair-end NextSeq sequencing, using the primers 515□F and 806□R which targets the V4 region of the 16S rRNA gene [38]. Samples obtained from DNA extraction were also amplified and quantified using qPCR (CFX Opus 96; Thermo Fisher, USA) for the bacterial and fungal DNA presence, as well as their ratio (B: F). Per 3 µl sample template, 1 µl of forward and reverse primer were added, along with 5µl of sterilized Milli-Q water and 10 µl of iQTM SYBR® Green Supermix (Bio-Rad, Hercules, USA). For bacteria, the primer pair was 515□F and 806□R which targets the V4 region of the 16S rRNA gene [38], while for fungi we used the primer combination FF390.1 and FR1 targeting the V7-V8 region of the 18S rRNA gene [39]. The PCR conditions were as follows: Denaturalization at 94 □C for 3 minutes, followed by 35 cycles of 95 °C for 15 seconds, 56 □C for 10 seconds, and an extension step at 72 □C for 30 seconds. Then, the bacterial and fungal DNA proportions in the soil were calculated from a standard linear regression curve obtained for each primer pair by making a dilution series of a solution containing a known concentration of DNA isolated from the soil and measuring the respective cycle threshold (Ct) values. For bacteria, this formula was Y = −3.62x + 16.58. For fungi, this formula was Y = −3.90x + 18.95. The Ct cut-off was set at 50 Relative Fluorescence Units (RFU) to filter out signals that were too weak and to identify signals that came up too late (Ct >30).

### 8. Microbial biomass and soil nutrient content determination

Microbial biomass C (MBC) and N (MBN) were determined using fumigation-extraction [40, 41]. MBC and MBN were measured using fumigation extraction, carried out in desiccators for 24 h at room temperature. Fumigated and non-fumigated samples of 10 g moist soil were mixed with 0.5 M K_2_SO_4_. Extraction was performed at 200 rpm, for 30 minutes. A multi-N/C 2100S analyzer was used to measure organic carbon (SOC) and nitrogen contents in the extracts. Microbial biomass C was calculated using the formula: (Extracted organic C from fumigated soil - Extracted organic C from non-fumigated soil) / 0.45 (correction factor) for MBC. MBN was calculated similarly: (Extracted organic N from fumigated soils - Extracted N from non-fumigated soil) / 0.54 (correction factor) for MBN. Results were used to determine organic carbon, total nitrogen, microbial biomass C (MBC) and microbial biomass N (MBN) concentrations.

### 9. Statistical analysis

#### 9.1 Plant parameters

All statistical analyses were performed in R software ver. 4.2.2. [42]. Figures were created using the package ggplot2 [43] and finalized using Adobe Illustrator ver. 27.9.4 (https://www.adobe.com/products/illustrator.html). Before the analyses of treatment effects, we checked for the residual normality and homoscedasticity of the models by visually inspecting the model assumptions using the ‘plot’ function. To test for the effect of predator addition on plant performance we constructed one mixed linear model for each plant parameter measured using the ‘nlme’ package [44], with ‘predator addition’, ‘nitrogen addition’ and their interactions fixed factors, and the ‘microbiome predator community’ and ‘microbiome predator diversity’ as random intercepts. The term ‘microbiome predator community’ refers to the 40 predator communities added to the experiment and ‘microbiome predator diversity’ refers to the diversity levels used. We then examined the significance using ANOVA for the different models created for each plant parameter.

To test the effect of both an increasing diversity of predators and an increasing N addition as well as their interaction on plant performance, we modelled both treatments as numerical variables. We again created a mixed linear model for each plant parameter measured using the ‘nlme’ package [44] and adding ‘microbiome predator community’ as a random effect factor. To account for increases in response variance to increasing levels of microbiome predator diversity and nitrogen addition, we allowed the model to estimate variance associated with our explanatory variables ‘microbiome predator diversity’ and ‘nitrogen addition’ using the ‘varExp’ function.

#### 9.2 Bacterial community analyses

The raw data obtained from the bacterial sequencing was processed using Qiime2 to remove chimeras, quality filtering, and finally obtain an ASV table. We used the SILVA database for taxonomic annotation of ASVs. The removal of primers and demultiplexing was performed by Genome Quebec (Quebec, Canada). We managed significant variations in read counts by excluding samples that deviated from the average read count by more than ± 2 standard deviations, which led to the removal of six samples with less than 12122 reads and one sample with more than 37433 reads. Following this, we eliminated all ASVs that appeared in fewer than four samples and had a maximum relative abundance below 0.01% to minimize the impact of rare ASVs on the community composition.

Following Deng et al [45] we only measured Shannon diversity as a proxy for alpha diversity for which we used the ‘alphaDiversity’ function from the ‘vegan’ package [46]. NMDSs were performed using the function ‘metaMDS’ from the ‘vegan’ package [46], using the following parameters: k = 3, distance = “bray”, trace = FALSE, autotransform = FALSE.

#### 9.3 Functional annotation

To detect how changes in bacterial community linked to changes in the functionality of the system, a functional annotation of the bacterial community was performed by using the ‘microeco’ package [47] and using the FAPROTAX database [48]. The bacterial community was first rarified using the lowest sum of reads per sample, which was 13695. This process resulted in the removal of 3 ASVs. Then, the functions were filtered in two ways: First, the ones with less than a 0.1 % of relative abundance were removed of the database. Secondly, we only selected the functions that light be linked to our experimental setup, ending up with the following functions: ‘Aerobic chemoheterotrophy’, ‘anaerobic chemoheterotrophy’, ‘cellulolysis’, ‘dark hydrogen oxidation’, ‘dark oxidation of sulfur compounds’, ‘fermentation’, ‘hydrocarbon degradation’, ‘intracellular parasites’, ‘nitrogen respiration’, ‘predatory or exoparasitic’, ‘respiration of sulfur compounds’, ‘ureolysis’.

Once function annotation was done and filtered, normality was assessed using the Shapiro-Wilk test. If data met the assumptions of normality and homogeneity of variances, parametric tests were used: Student’s t-test for two-level comparisons (*PredatorsAdded*) and ANOVA for multi-level factors (*NitrogenAdd* and *SpeciesNumber*). If assumptions were violated, non-parametric alternatives were applied: Wilcoxon rank-sum test or Kruskal-Wallis test, respectively.

Additionally, effect sizes were estimated by extracting R² values from ANOVA models as a measure of explanatory power or, in cases where normality was not met, a pseudo-R² (□^2^) was estimated as analogous to R².

#### 9.4 Differential Abundance Analyses

In order to identify some potential genus of bacteria that might be influencing the response of the plant biomass, we performed differential abundance analyses using Deseq2 package in R [49]. Additionally, we only showed the genera that were statistically significant (p < 0.05) and visually highlighted only genera with a log2 median ratio bigger than +-2.The differential abundance analyses comparisons were done among predator addition (Yes vs No) and predator diversity (all the pair combinations between treatments of 0, 6, 12 and 24 species). The results of those analyses are depicted in the figures (P > 0.05 = ns; P < 0.05 = *; P < 0.01 = **; P < 0.001 = ***).

#### 9.5 Correlation analyses

To obtain correlations between all the variables and parameters measured we used Pearson correlations with the function ‘rcorr’ from the ‘Hmisc’ package [50]. Prior to the correlations, binary categorical variables (e.g., “Yes“/“No”) were converted to numeric (0/1) format for inclusion in the correlation matrix. To correct for multiple testing across all pairwise correlations, False Discovery Rate (FDR) correction was applied using the Benjamini-Hochberg procedure. A corrected p-value threshold of 0.05 was used to assess significance. For the correlation matrix plot, we used the ‘corrplot’ package [51].

#### 9.6 Structural Equation Modeling (SEM)

To further examine how independent and interactive impacts of nitrogen addition and microbiome predator diversity directly or indirectly, through changes in microbial communities, ultimately affected plant biomass we constructed a structural equation model using the piecewiseSEM package[52]. We hypothesized that N addition, microbiome predator diversity and their interaction could directly affect microbial N (mg/kg) and bacterial community composition (first NMDS axis). Within the soil microbial community, we hypothesized that bacterial community composition affected microbial N (mg/kg). Finally, we hypothesized that total plant biomass may be driven by bacterial community composition, changes in microbial N, as well as by the direct effect of nitrogen addition. In this model, we natural-log-transformed total plant biomass, microbial N (mg/kg), nitrogen addition (Ln + 1) and microbiome predator diversity (Ln + 1). Microbiome predator community was removed as a random effect after its initial addition, as it did not explain variation

## Results

### The addition of microbiome predators affects plant growth but not nutrient content

Overall, microbiome predator addition, independent of diversity levels, increased total plant biomass by 53% (X^2^ = 28.34 ; df = 1; P < 0.001; Figure 2A). This positive effect on plant growth was shown for both shoot (52%, X^2^ = 13.74 ; df = 1; P < 0.001; Supplementary Figure 1A) and root biomass (66%, X^2^ = 21.2 ; df = 1; P < 0.001; Supplementary Figure 1B). The root-shoot ratio was not affected by any of the treatments (X^2^ = 0.45 ; df = 1; p-value = 0.5; Supplementary Figure 1C). Opposite to plant biomass, microbiome predator addition did not affect plant nutrient content parameters (Chlorophyll content, leaf C, leaf N, leaf S and leaf C/N ratio), (P > 0.05; Figure 2).

**Figure 2.**
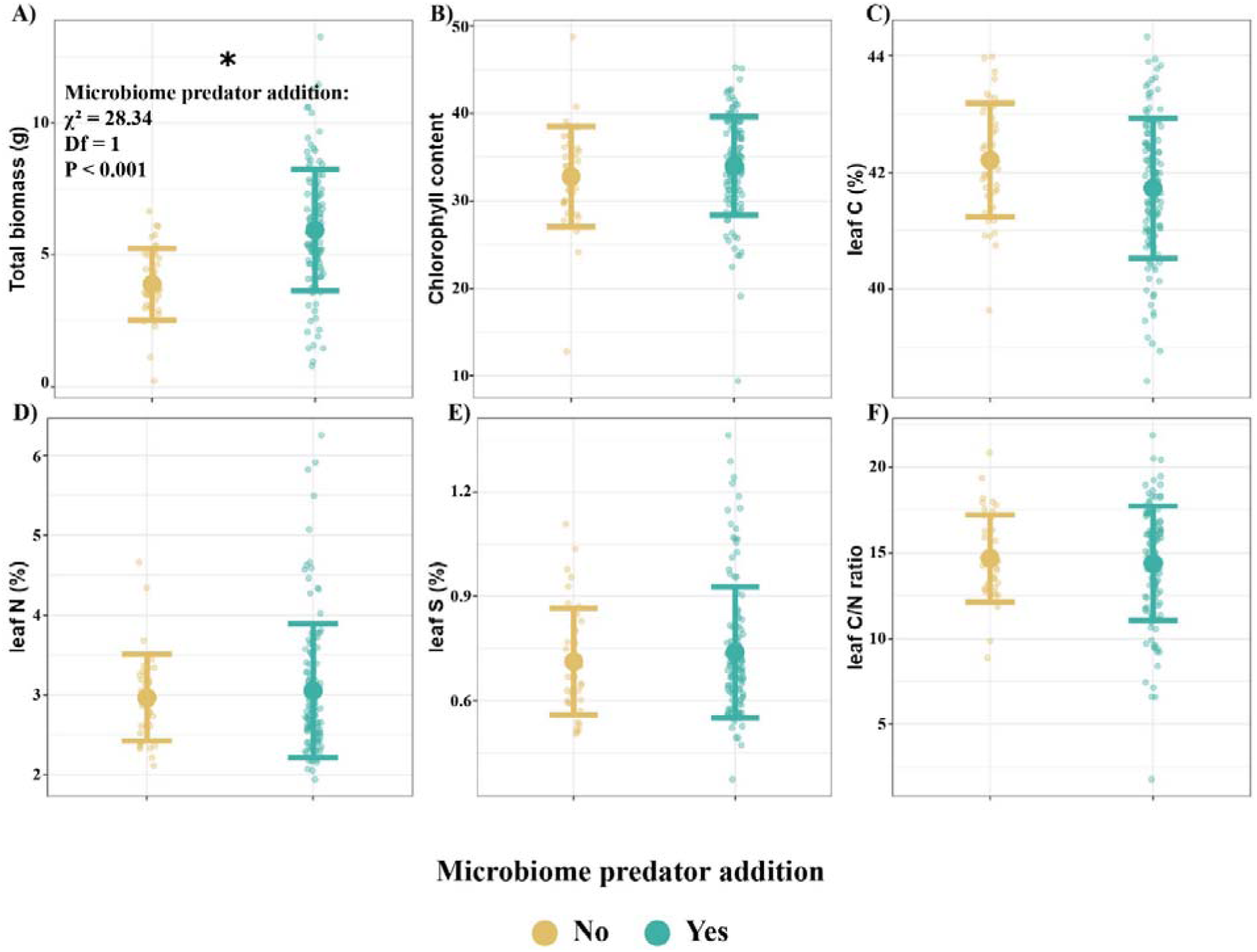
Microbiome predator addition increases total plant biomass (g) (A) but not chlorophyll content (B), leaf C (C), leaf N (D), leaf S (E), leaf C/N ratio (F) irrespective of diversity levels (X-axis shared for all panels). Small dots represent the raw data, error bars represent the standard deviation, and the thicker dot represents the mean. Statistically significant effects (ANOVA; P < 0.05) are shown with an asterisk.

The positive effect of microbiome predator addition, irrespective of diversity levels, on plant biomass was not related to the amount of N added (P = 0.61; X^2^ = 1.81, df =1 ; Supplementary Figure 2).

### Total plant biomass depends on increasing diversity of microbiome predators and nitrogen levels

To understand relationship between microbiome predator diversity and plant performance we analyzed the impact of increasing diversity of microbiome predators (0 - 20 protist species; 0 - 4 nematode species) on different plant parameters related to plant biomass and leaf nutrient content. We found that both an increasing diversity of microbiome predators and an increasing addition of nitrogen separately had a positive effect on plant biomass and leaf chlorophyll and nutrient contents (Figure 3; P < 0.05), while no significant interactions between both independent variables were found (ANOVA; all P-values > 0.05). The only response variables that were not significantly affected by increasing microbiome predator diversity were leaf C (%) (Fig 3E), leaf N (%) (Fig 3G) and leaf C/N ratio (Fig 3K). From the other variables, plant biomass most strongly benefited with increasing microbiome predator diversity (X^2^ = 13.91; P < 0.001; d.f. = 1), while chlorophyll content was most strongly affected by N levels (X^2^ = 13.69; P < 0.001; d.f. = 1). Except for total plant biomass, increasing nitrogen levels (as shown by P and X^2^) affected plant parameters more strongly, than microbiome predator diversity.

**Figure 3.**
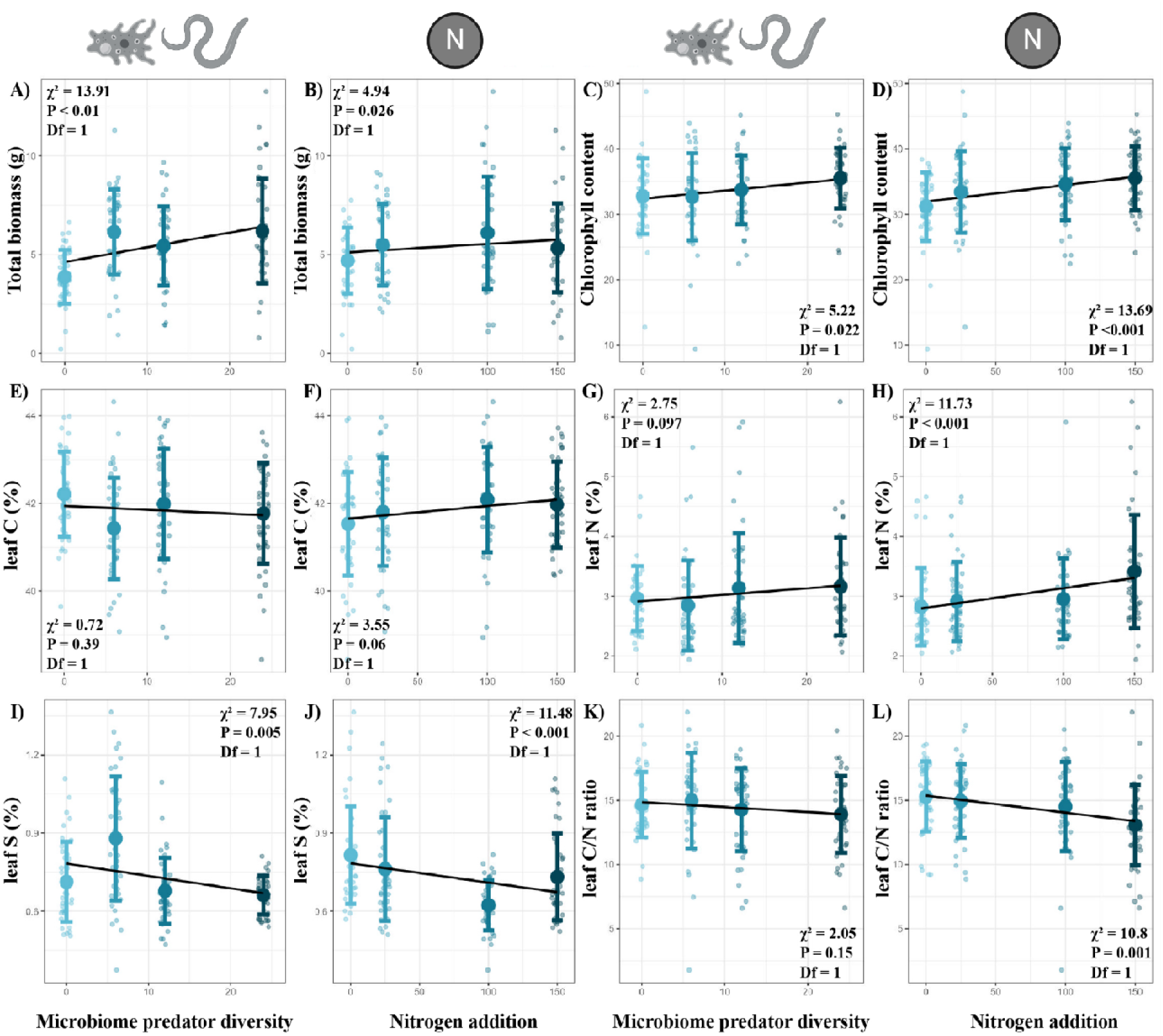
The relationship between plant total biomass (g) (A, B), Chlorophyll content (C, D), leaf C (%) (E, F), leaf N (%) (G, H), leaf S (%) (I, J), and leaf C/N ratio (K, L) with an increasing diversity of microbiome predators (A, C, E, G, I. K) or an increasing N addition (B, D, F, H, J, L). Small dots represent the raw data, error bars represent the standard deviation, and the thicker dot represents the mean. The test results shown in the panels are based on the linear mixed-effect models performed for each plant parameter. The X-axis is shared per column.

While no interactive effect of microbiome predator diversity and N addition on plant biomass was found, they interactively affected plant biomass when N addition was modelled as a factor (ANOVA; P < 0.001; X^2^ = 18.07; Supplementary Figure 3).

### Microbiome predators rather than N addition shaped bacterial community composition

We analyzed the bacterial community composition as one of the potential mechanisms behind the impact of predator and N addition, as well as microbiome predator diversity and N levels, on plant performance. The bacterial community composition was mostly affected by the sole addition of microbiome predators (PERMANOVA; P < 0.001; F-value = 15.5; R^2^ = 0.09; Figure 4A) but also by N addition (PERMANOVA; P < 0.001; F-value = 2.4; R^2^ = 0.017; Figure 4B). Moreover, the bacterial community composition was impacted by microbiome predator diversity (PERMANOVA; P < 0.001; F-value = 8.99; R^2^ = 0.15; Figure 4C) and N levels (PERMANOVA; P < 0.001; F-value = 2.39; R^2^ = 0.046; Figure 4D).

**Figure 4.**
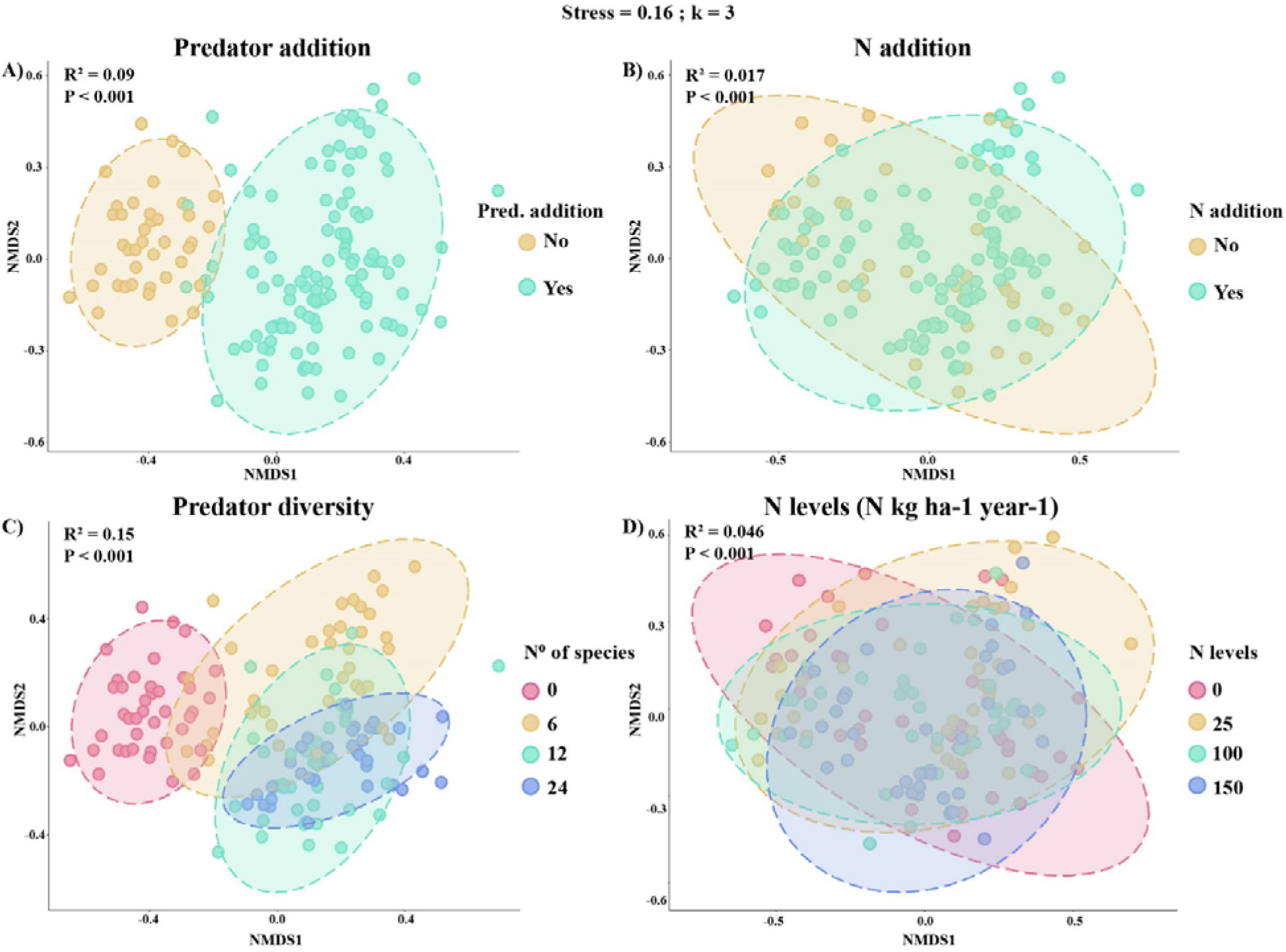
NMDSs representing the bacterial community composition at the ASV level depending on predators (A) or N (B) addition, as well as on an increasing diversity of predators (C), or an increasing level of N added (D), showing that microbiome predators rather than N are key players in shaping bacterial communities. The test results shown in each of the panels refer to PERMANOVA analyses. Stress and k values, indicating the overall fit of the data and the number of NMDS-axes used within the ordination space, respectively, are indicated.

While bacterial community composition was mainly shaped by microbiome predator addition, alpha diversity (Shannon index) did not change with the addition of microbiome predators (P = 0.56; Supp Fig 4A) or nitrogen (ANOVA; P = 0.89; Supp Fig 4B). However, alpha diversity did change with the interaction between predator diversity and nitrogen addition (ANOVA; Χ^2^ = 8.11; Df = 1; P < 0.01; Fig 5A-D).

**Figure 5.**
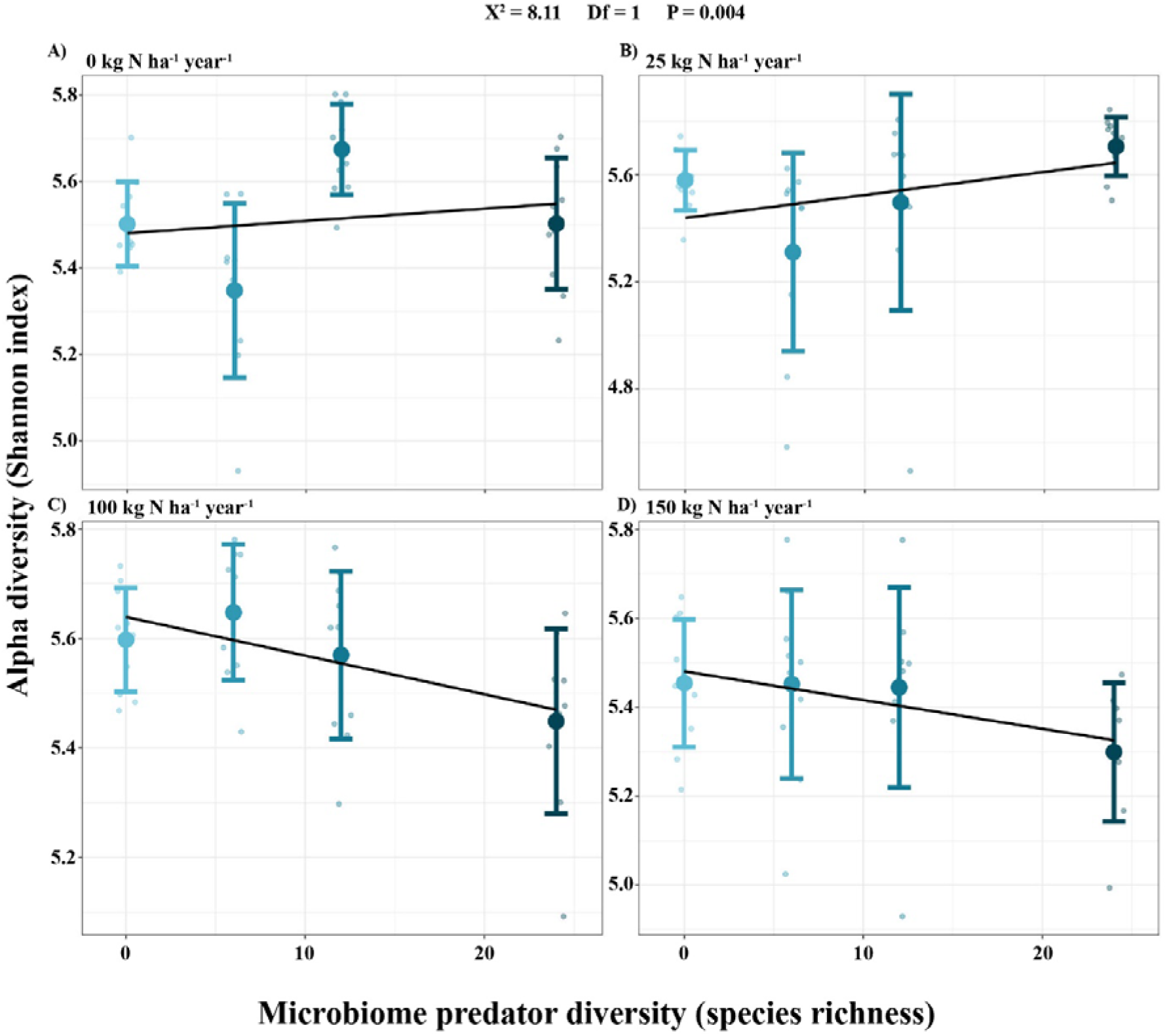
Relationship between Shannon (A-D) and chao1 (E-H) alpha diversity indexes varies with an increasing microbiome predator addition (X-axis) per level of nitrogen addition. The test results shown above the figure are based on the linear mixed-effect models performed to test the interaction between increasing diversity and N levels. The nitrogen levels are written above of each subplot.

### Differential abundance analyses

To detect bacterial groups that most strongly linked with changes in plant performance we performed differential abundance analyses. Microbiome predator addition increased the abundance (log2 fold change > 2) of 30 genera, while it decreased the abundance (log2 fold change < −2) of 7 genera (Supplementary Figure 5). The three genera that increased the most were *Alicyclobacillus* (6.76 log2 fold change), *Filibacter* (5.61 log2 fold change) and *Rugamonas* (5.57 log2 fold change). The ones that decreased the most were *Daeguia* (−2.17 log2 fold change), *Leptospira* (−2.54 log2 fold change), and *Paludibaculum* (−3.14 log2 fold change).

When comparing treatments with different levels of predator diversity, the largest number of genera showing log2 fold changes > 2 or < −-2 occurred in the comparisons between the predator addition treatments (6, 12, or 24 species) and the control treatment with no predators (0 species), while also several taxa were differentially abundant between the diversity levels (Supplementary Figure 6).

### Functional annotation of bacterial communities

To determine whether the changes in bacterial composition were linked to potential changes in potential bacterial-driven functions, we functionally annotated the bacterial taxa (using FAPTROTAX [48]) and focused on 12 functions related to plant performance as well as N and C cycling. Among those 12, microbiome predator addition significantly affected 8 of them (Figure 6), as it increased ‘aerobic chemoheterotrophy’ (11.51%, Wilcoxon; P < 0.001), ‘predatory or exoparasitic’ (359%, Wilcoxon; P < 0.001) and ‘hydrocarbon degradation’ (350.6%; Wilcoxon; P < 0.001), while reducing ‘cellulolysis’ (−38.5%, Wilcoxon; P < 0.001) ‘dark oxidation of sulfur compounds’ (−18.9%, Wilcoxon; P < 0.001), ‘fermentation’ (−15%; Wilcoxon; P < 0.01), ‘nitrogen respiration’ (−28%, T-test; P < 0.001) and ‘ureolysis’ (−43%, Wilcoxon; P < 0.001).

**Figure 6.**
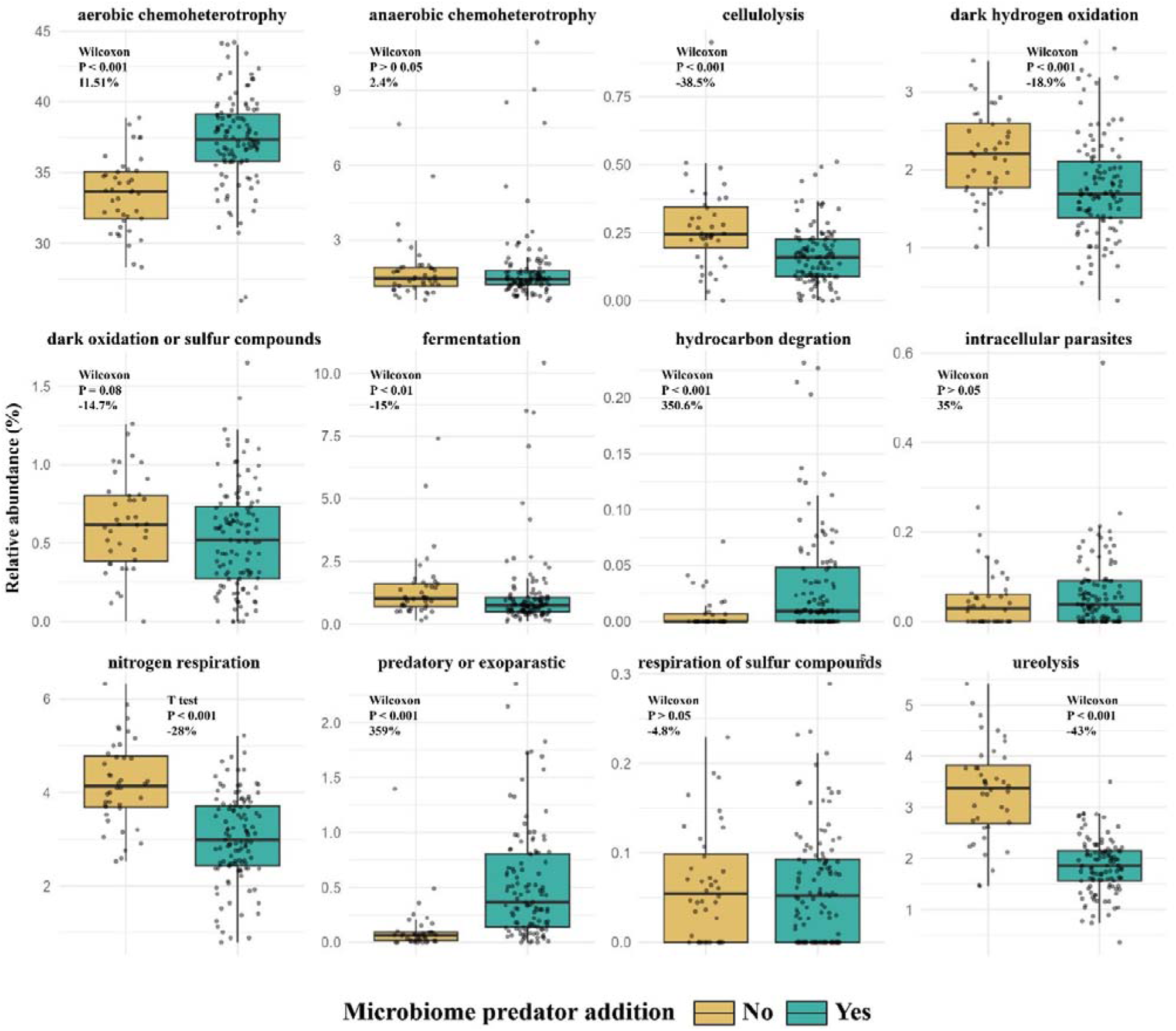
Boxplot depicting the effect of predator addition on the relative abundance of the different bacterial functions. Statistical tests performed, together with the p value and the percentage of relative abundance changed are shown within each panel.

When testing the effect of microbiome predator diversity on the same 12 functions, only respiration of sulfur compounds was not affected (Figure 7). ‘Aerobic chemoheterotrophy’ (R^2^ = 0.28; ANOVA; P < 0.001), ‘hydrocarbon degradation’ (□^2^ = 0.17, Kruskal-Wallis; P < 0.001), ‘intracellular parasites’ (□^2^ = 0.08, Kruskal-Wallis; P = 0.001) and ‘predatory or exoparasitic’ (□^2^ = 0.27, Kruskal-Wallis; P < 0.001) functions were enhanced compared to control, while ‘cellulolysis’ (□^2^ = 0.1, Kruskal-Wallis; P < 0.001), ‘dark hydrogen oxidation’ (R^2^ = 0.14; ANOVA; P < 0.001), ‘fermentation’ (□^2^ = 0.08, Kruskal-Wallis; P < 0.001), ‘nitrogen respiration’ (R^2^ = 0.27; ANOVA; P < 0.001) and ‘ureolysis’ (□^2^ = 0.38, Kruskal-Wallis; P < 0.001) decreased.

**Figure 7.**
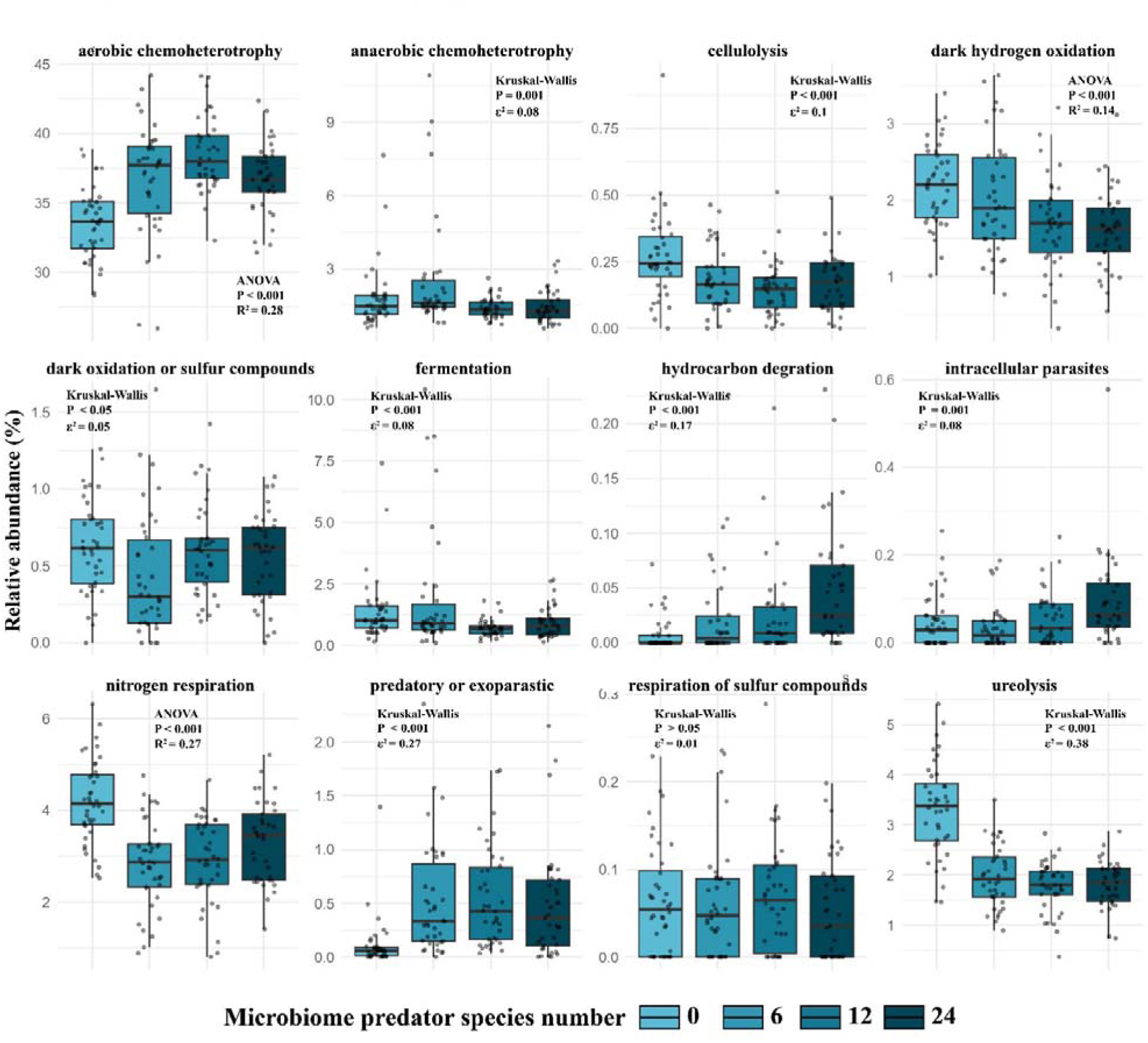
Boxplot depicting the effect of increasing protists diversity on the relative abundance of the different bacterial functions. Statistical tests performed, together with the p value and the R^2^ or a pseudo-R^2^ (□) are shown within each panel.

### SEM

To statistically test the potential mechanisms through which microbiome predator diversity and N levels affected plant biomass, we constructed an SEM (Fig. 8). As we acknowledge the hypothetical effect of some other variables not used in the SEM, we performed a correlation analysis with all the variables measured in the study to visualize an overall picture (Supplementary Figure 7). Our structural equation model presented a good fit (P = 0.158; Fisher’s C = 3.692; 2 d.f), showing that N addition and microbiome predator diversity, solely and interactively, affected total plant biomass through impacts on microbial N and bacterial community composition. Both N addition (Std. estimate = 0.68; P < 0.001) and microbiome predator diversity (Std. estimate = 0.38; P = 0.054) separately had a positive effect on MBN (Microbial Biomass Nitrogen), but the interaction of both negatively affected MBN (Std. estimate = −0.41; P = 0.03). These changes in MBN did not result in changes in plant biomass, but N addition directly increased plant biomass (Std. estimate = 0.18; P = 0.02). N addition, microbiome predator diversity and the combination of both treatments significantly affected the bacterial community composition. Among these three, microbiome predator diversity was the main factor determining bacterial community composition (Std. estimate = 0.98; P <0.001), which was the main variable affecting plant biomass (Std. estimate = 0.49; P <0.001), (Fig. 8).

**Figure 8.**
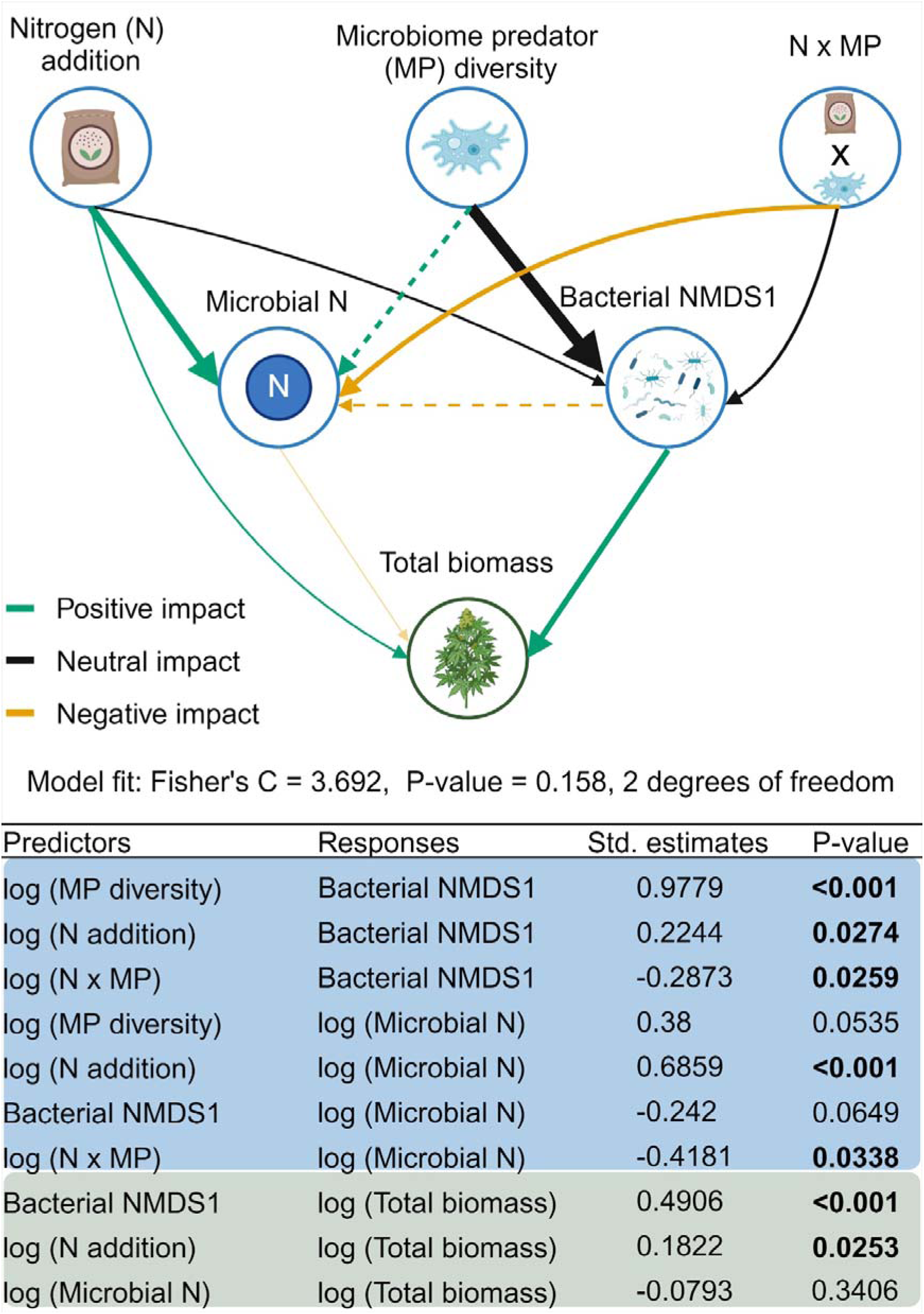
In the upper part the representation of the SEM model is visualised, in which it is observed that plant biomass is most affected by bacterial community composition, which is most affected by microbiome predator diversity. Arrow widths are determined by their standardized estimates, while arrow colors depict the type of relationship (positive, negative or neutral). “Neutral” denotes types of relationship with non-definable directional changes. Dashed lines represent marginally significant paths (0.05 < p value < 0.07). The bottom part shows the statistics of the model for each path. The colors of the table are linked to the colors on the circles surrounding each icon in the scheme above.

## Discussion

Our study demonstrates that increasing soil microbiome predator diversity can enhance plant biomass without affecting plant nutrient content, while nitrogen addition predominantly improves plant nutrient content. Overall, microbiome predator addition increased plant biomass by 53%. We further found that microbiome predator diversity was key in shaping bacterial community composition and, consequently, potential ecosystem functions related to N cycling as well as plant growth.

### Microbiome predator addition on plant performance

Microbiome predator addition, irrespective of diversity levels, increased plant biomass by about half without affecting plant nutrient contents. These findings are partially in line with our first hypothesis and previous studies based on bacterivorous protists or nematodes [53–55]. Although the positive effect of microbiome predator addition on plant biomass was observed across all nitrogen (N) levels, the effect appeared more pronounced under low and medium N addition. This trend may suggest that N limitation was alleviated in the highest N treatments, reducing the relative benefit of predator addition. However, given the marginal statistical significance of these results (see Supplementary Figure 2), it is possible that other nutrients—such as phosphorus (P) and potassium (K)—contributed in determining plant biomass. These nutrients have also been shown to be released by microbiome predators through predation on the microbiome [23, 56]. While biomass increased, plant nutrient content (chlorophyll content, leaf C, leaf N, leaf S and leaf C/N ratio) was not affected by microbiome predator addition. This result contradicts our first hypothesis, but is supported by research showing that an increase in plant biomass due to protist addition is not necessarily linked to an increase in nutrient content in the shoots [57]. This apparent contradiction may be explained by the ability of microbiome predators to release essential nutrients beyond N and C—such as P and potassium (K) [56, 58], which may have contributed to more balanced plant nutrition. Apart from a more balanced plant nutrition, microbiome predators can also release certain growth hormones either directly or via changing the microbiome composition, increasing plant biomass without affecting the relative nutrient concentrations [23]. In contrast, N addition had minimal positive effects on biomass but notably improved plant nutrient content. This aligns with studies indicating that excess N is stored in the plant without affecting biomass when other nutrients (P,K) are a limiting factor instead of N [23, 56].

### Microbiome predator diversity on plant performance

Similarly to microbiome predator addition, increasing microbiome predator diversity led to a plant biomass increase, in line with our second hypothesis. Contrary to results of microbiome predator addition, their diversity interacted with N levels to determine plant biomass, with significant effects observed under none, low, and mid N levels, but not under high N conditions (Supplementary Fig. 3). This pattern aligns with Liebig’s law of the minimum, which states that plant growth is constrained by the most limiting resource [59]. Thus, at high N levels, where nitrogen was no longer limiting, the influence of microbiome predator diversity diminished. This microbiome predator diversity-induced increase in plant biomass aligns with the BEF concept, which states a positive relationship between diversity and ecosystem functioning [8]. This concept has been largely tested and documented aboveground, with plant diversity positively affecting plant productivity in famous experimental setups such as Cedar Creek or the Jena experiment (e.g. [60, 61]), but barely experimentally tested with soil biodiversity [14]. The main explanation behind the positive relationship is thought to be the complementarity of niches (time, space, feeding habits, etc) among different species. The niche complementarity, in our setup, is derived from species-specific predation differences among nematode and protist species [62–64], broadening the potential feeding niches with increasing species richness of predators. In this way, more nutrients might be released to be taken up by the plant. This reinforces the idea that sharing the same function (here microbiome predation) does not imply functional redundancy but complementarity [65]. However, our results also showed that increasing N levels, rather than microbiome predator diversity, had an impact on all plant nutrient content parameters, which is in line with other studies showing N-dependency responses of leaf nutrient content [66]. This plant nutrient N-dependency was shown to be independent of microbiome predator diversity, indicating that the benefits from enhancing microbiome predator diversity existed regardless of the N addition levels and vice versa.

### Microbiome predator diversity shaped plant biomass via bacterial community composition

Partially in contrast with our third hypothesis, increasing microbiome predator diversity was correlated with total plant biomass and bacterial community composition but not with soil nutrients or other soil and plant parameters. The clear distinction in our bacterial community analyses regarding microbiome predator diversity, and the strong link shown in the SEM analyses between microbiome predator diversity and bacterial community composition, emphasizes the feeding preferences of microbiome predators [62, 63, 67]. Thus, increasing microbiome predator diversity appears to enhance the complementarity of feeding niches, likely enabling a broader range of bacterial taxa to be targeted. These targeted feeding patterns are often associated with increases in plant growth-promoting bacteria or reductions in plant pathogenic bacteria, ultimately contributing to increased plant biomass [28, 53, 68–70]. In our case, while we showed that certain plant-beneficial genera such as *Burkholderia* (reported as plant growth-promoting [71]) or *Kaistia* (reported to suppress the fungal plant pathogen *Fusarium oxysporum* [72]) were enhanced in treatments with increasing microbiome predators, changes in the bacterial functional profile are of more importance. For example, N cycling functions like ‘nitrogen respiration’ and ‘ureolysis’ declined with increasing microbiome predator diversity, suggesting reduced N losses to the environment and greater N retention for plant use. C cycling was also enhanced as ‘aerobic chemoheterotrophic’ bacteria, which are linked to the formation of organic matter in soils [73], increased with an increasing protist diversity. We thus support the fact that microbial loop involves more processes than just N release, as demonstrated by Clarholm [25] and Bonkowski [23]. Additionally, we reveal the profound changes in the functional profile of the bacterial communities induced by predation. Finally, our results show that the effects of microbiome predator diversity on the microbial and auxiliar loops also apply when N is added. Consequently, the sBEF concept could be used as framework for agricultural systems unlike some other approaches, which only work in low nutrient conditions.

## Conclusions

In this way, we enhance the scientific body of evidence supporting the central role of microbes and their predators in nutrient cycling (i.e. microbial loop theory) by empirically linking microbiome predator diversity to plant performance. Furthermore, the data presented here recognize microbiome predators as potential biostimulants, as previously suggested [26].

In summary, we demonstrate that N addition and microbiome predator diversity may play different and complementary roles in plant performance: increasing N levels affected plant nutrient content, while microbiome predator diversity mainly affected plant biomass. Our findings show that microbiome predator diversity indirectly enhances plant biomass via changes in the soil bacterial community composition as well as by by promoting nutrient release, probably beyond only N. Finally, as microbiome predators caused the highest biomass in low or medium levels of N, we propose the potential use of microbiome predators as sustainable biostimulants with the aim to reduce the need for N inputs while maintaining or increasing plant biomass and nutrient content.

## Author contributions

A.B.G, S.G., J.B, N.H and F.W. conceived of the study and designed the experiments. J.B and N.H performed the experiment. J.B, N.H, A.B.G, S.G and F.W obtained all the data. A.B.G and R.W. analyzed the data. A.B.G and S.G. wrote the original manuscript and all coauthors contributed to its revision. All authors have read and approved the final version.

## Acknowledgements

Authors want to acknowledge Bram Willems for his help setting up the experiment as part of his BSc thesis.

