## Supplementary for "Soil microbiome predator diversity outperforms nitrogen addition in boosting plant biomass via bacterial community shifts"

**Supplementary material**

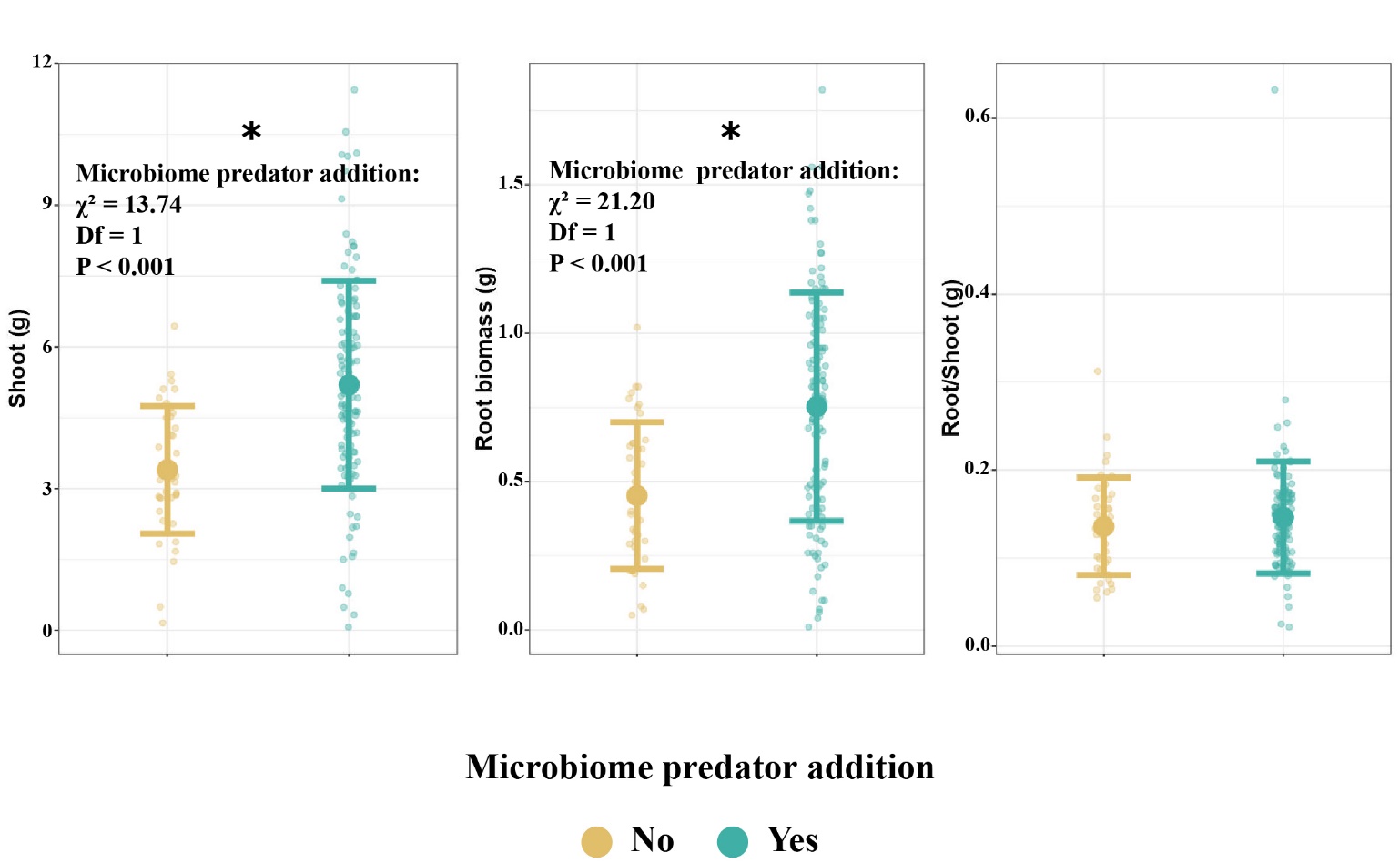

Supplementary Figure 1. Microbiome predator addition increases both shoot (A) and root (B) but not the root/shoot ratio (C) irrespective of diversity levels (X-axis shared for all panels). Small dots represent the raw data, error bars represent the standard deviation, and the thicker dot represents the mean. Statistically significant effects (ANOVA; P < 0.05) are shown with an asterisk.

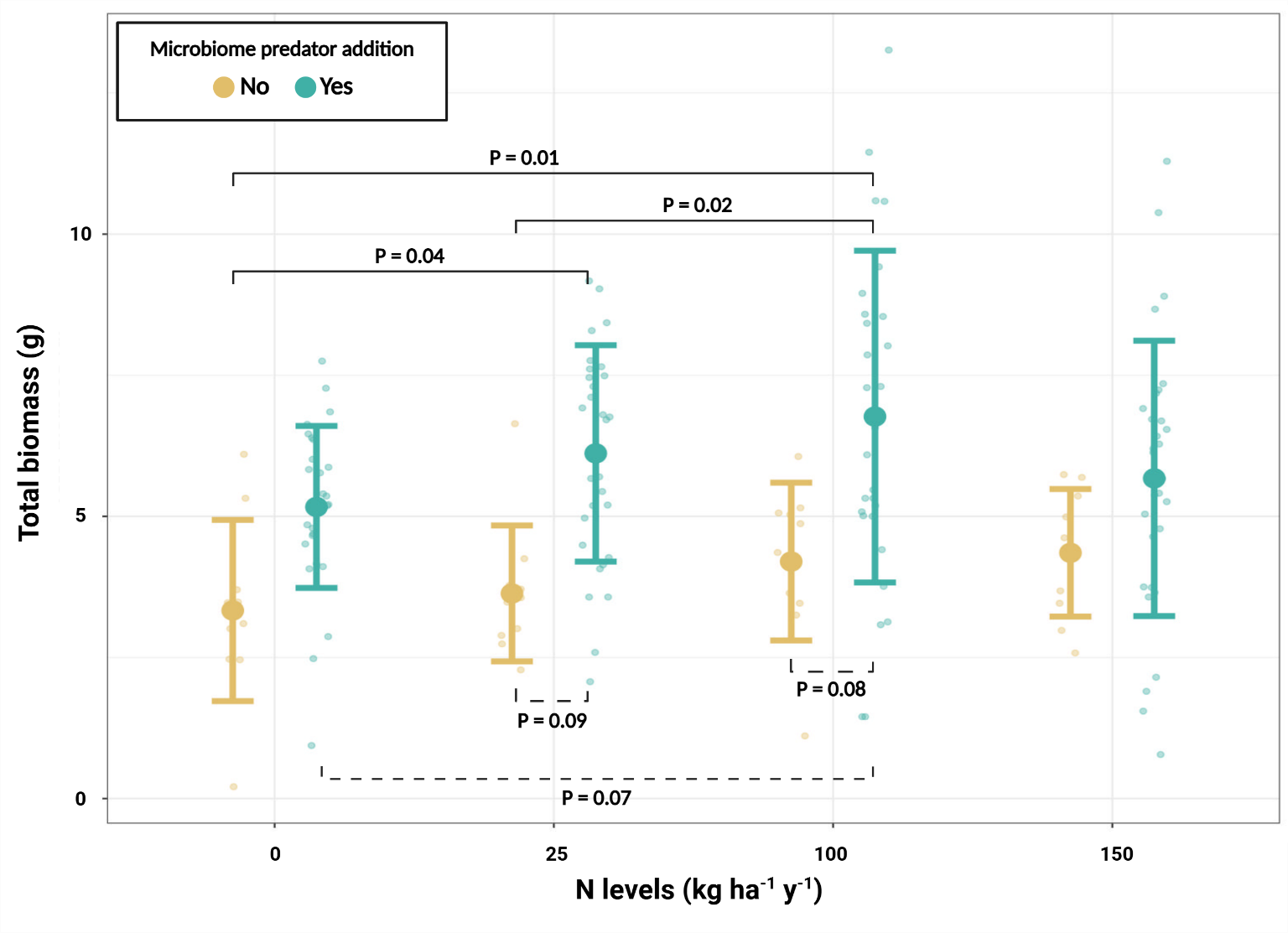

Supplementary Figure 2. Microbiome predator addition enhances plant total biomass (g) (Y-axis), reaching its maximum at low and mid N levels (X-axis). Small dots represent the raw data, error bars represent the standard deviation, and thicker dots represent the mean. Lines connecting treatments represent significant (continuous line) or marginally significant (0.05-0.1; dashed line) post hoc Tukey tests.

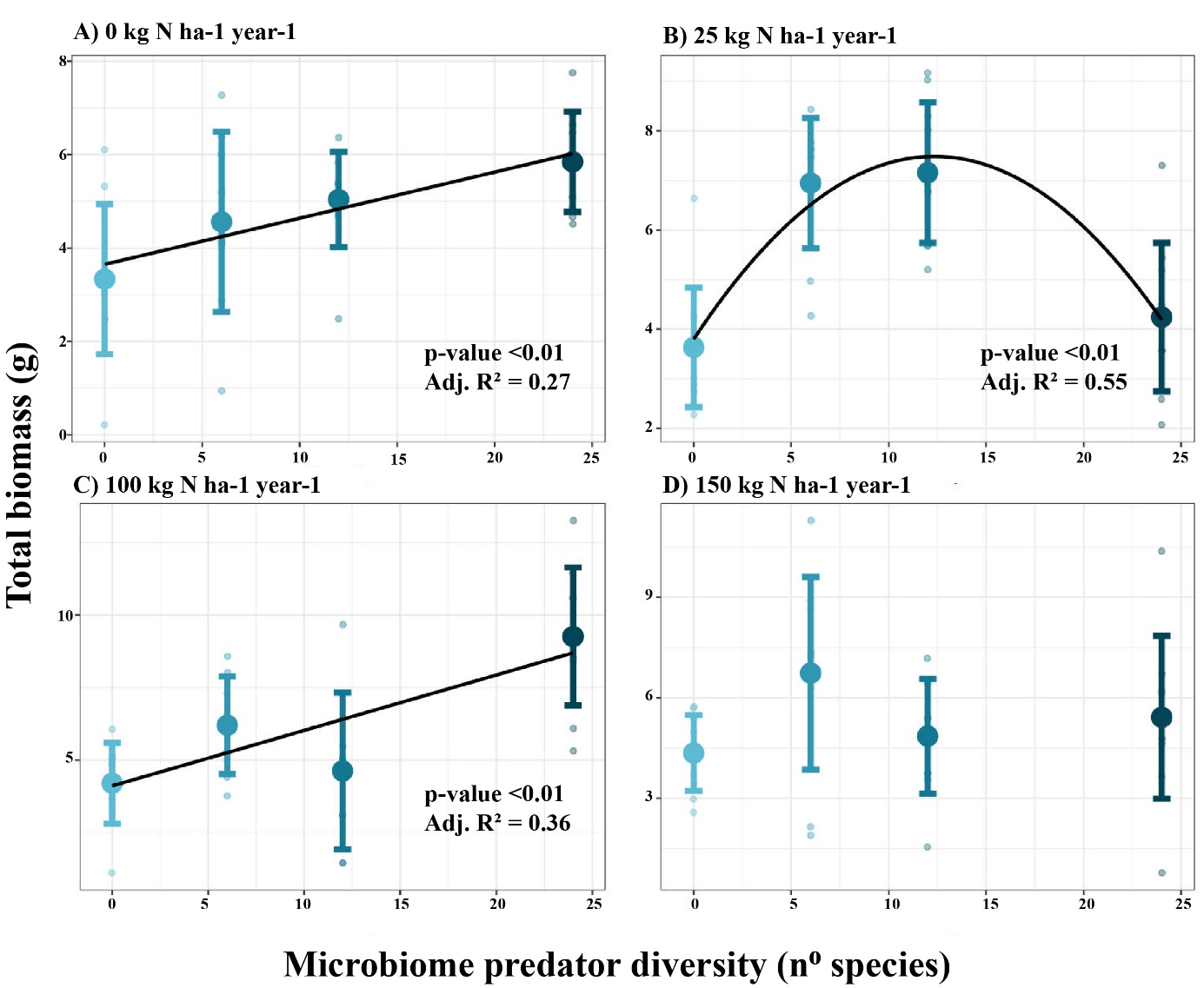

Supplementary Figure 3. The relationship between plant total biomass (g) and increasing diversity of microbiome predators is dependent on the different N levels (A, B, C, D). Small dots represent the raw data, error bars represent the standard deviation, and the thicker dot represents the mean. The test results shown in the panels are based on the individual models performed for each N level. The X-axis is shared per column.

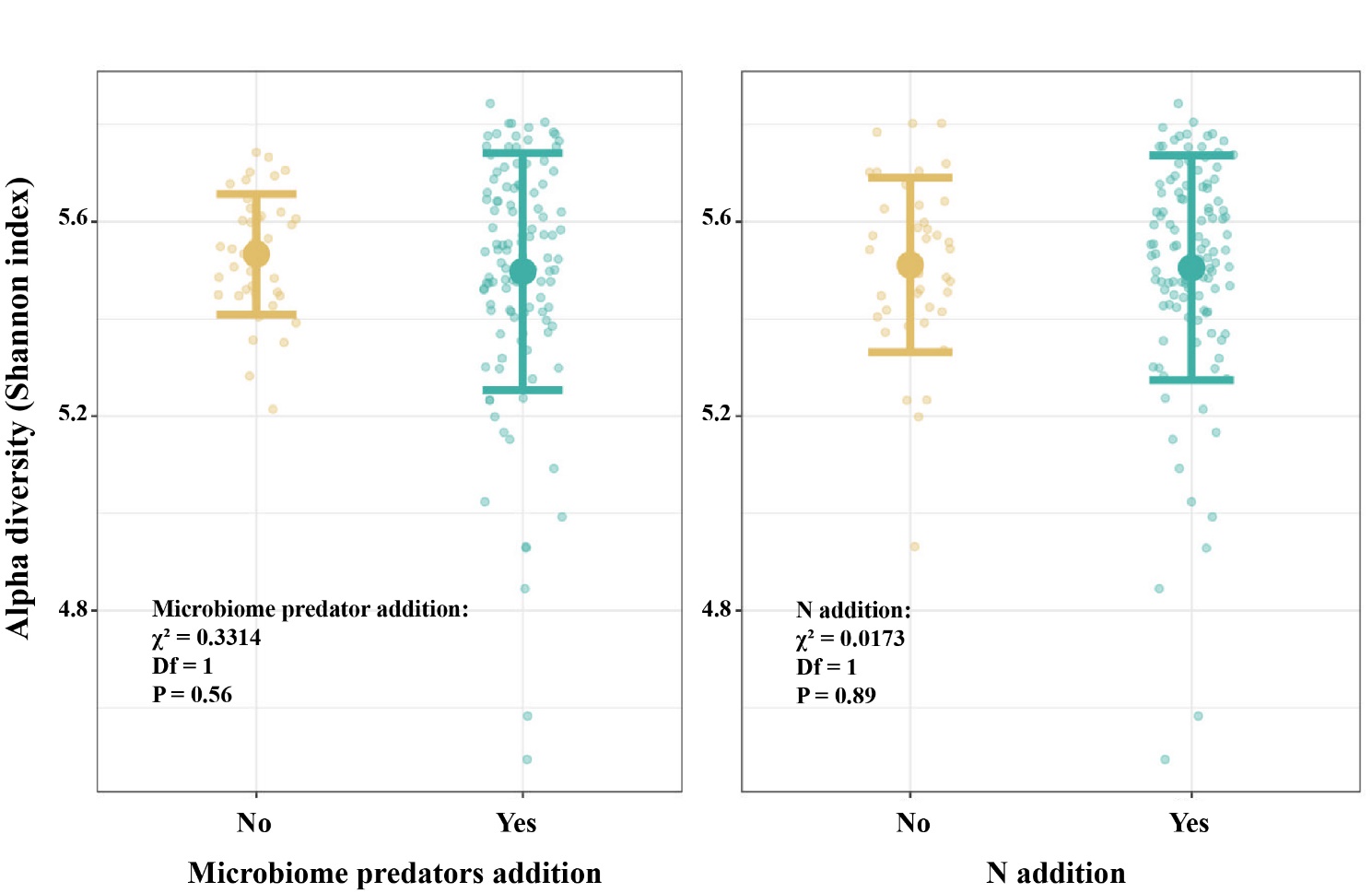

Supplementary Figure 4. Relationship between Shannon alpha diversity index with microbiome predator (A) and nitrogen addition (B). Small dots represent the raw data, error bars represent the standard deviation and the thicker dot represents the mean.

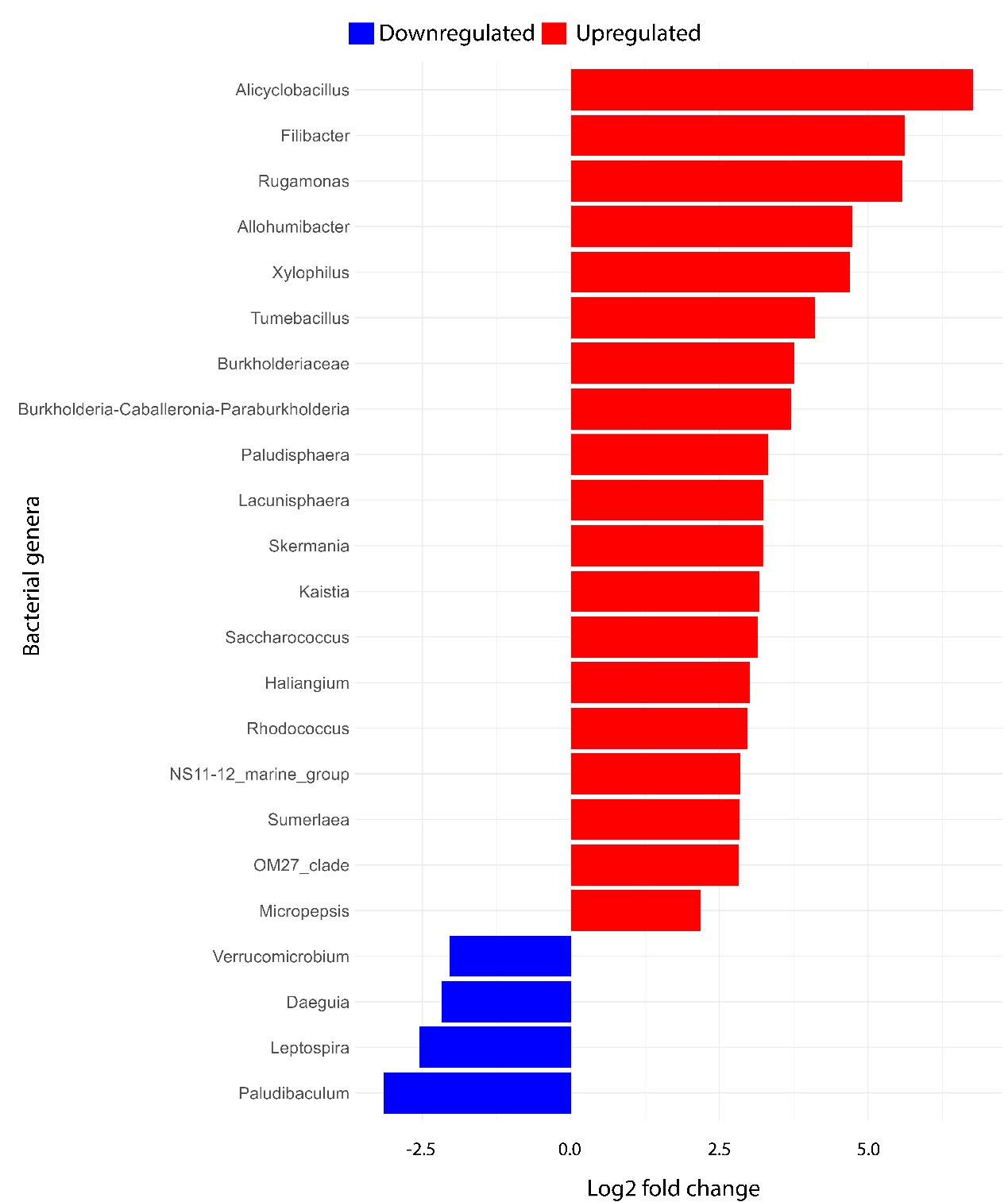

Supplementary Figure 5. Differential abundance analysis of bacteria at genus level. Treatments compared are microbiome predator addition versus not addition. Red bars on top and blue bars on the bottom represent statistically significant genera with a Log2 fold change higher than 2 and lower than -2 respectively. Genera with values >-2 and <2 where removed for readability purposes. The results should be seen as follows: positive Log2 fold change values: those genera increased when microbiome predators were added. Negative Log2 fold change values: those genera decreased when microbiome predators were added.

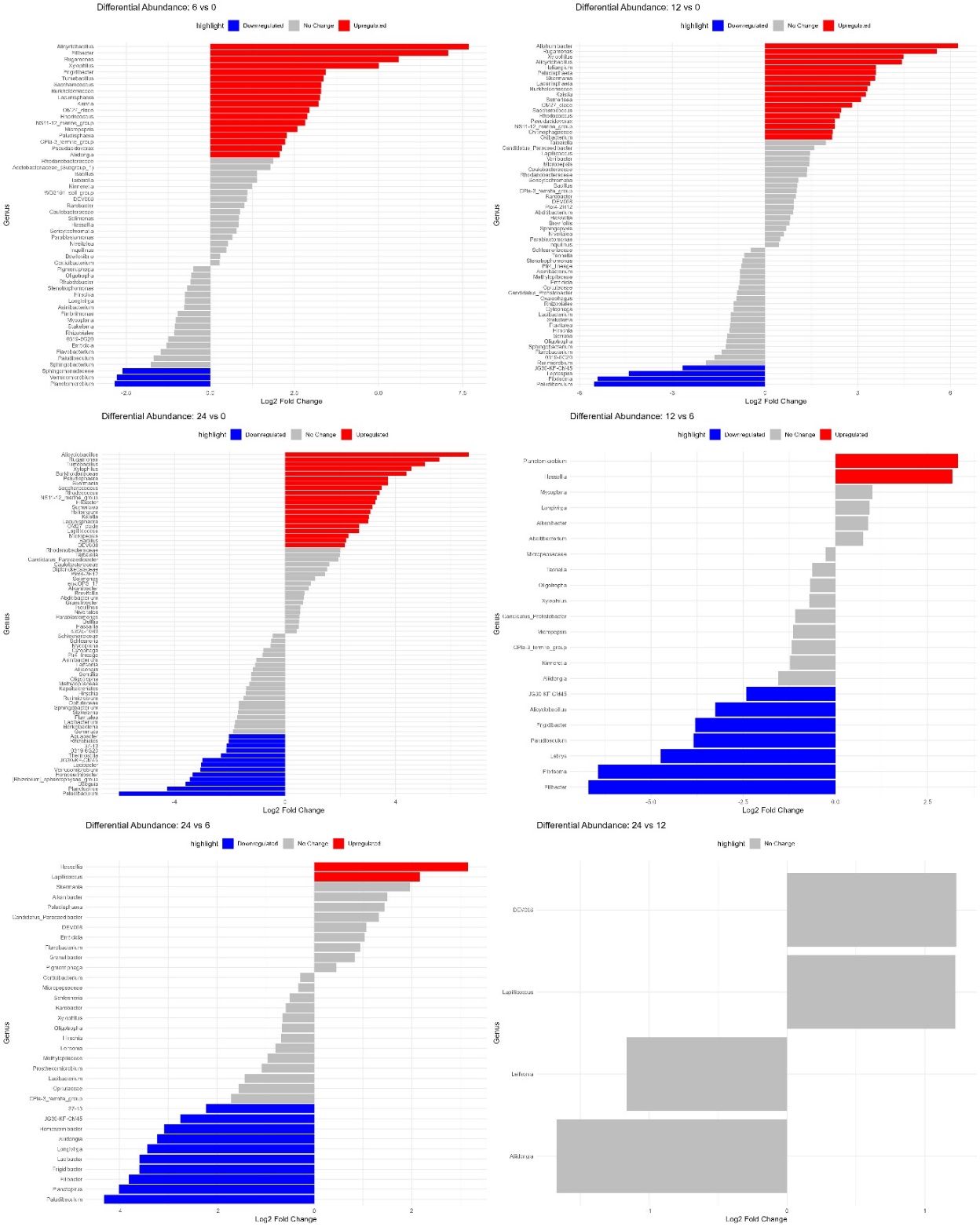

Supplementary Figure 6. Differential abundance analysis of bacteria at genus level. Treatments compared are shown on top of each graph. From left to right: 6 vs 0 species, 12 vs 0 species, 24 vs 0 species, 12 vs 6 species, 24 vs 6 species and 24 vs 12 species. Red bars on top and blue bars on the bottom represent statistically significant genera with a Log2 fold change higher than 2 and lower than -2 respectively. These results must be understood as genera that either increased or decreased in relative abundance when comparing the first treatment against the second.

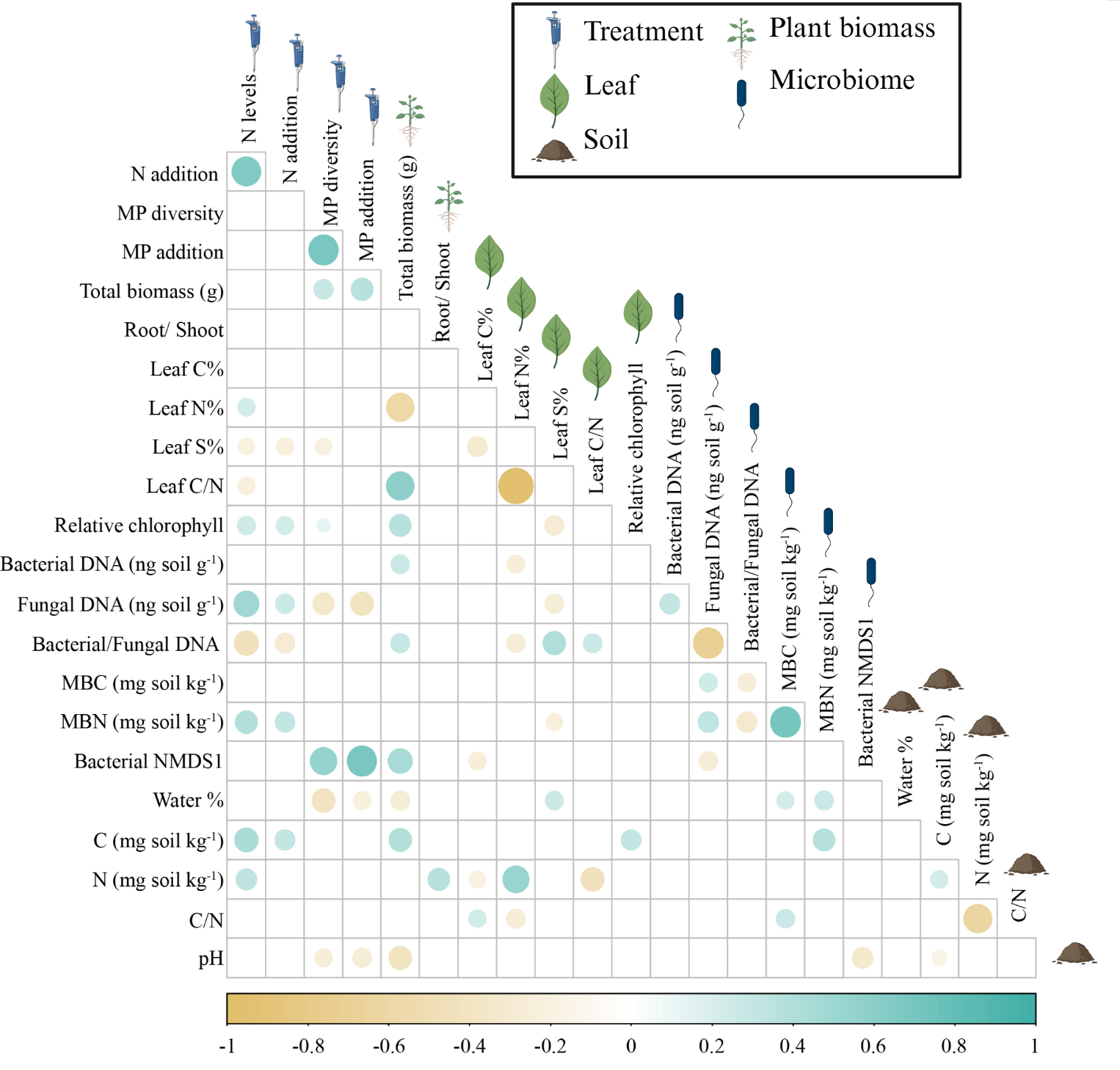

Supplementary Figure 7. Correlation plot among biotic and abiotic parameters measured. Empty spots represent no statistical significance (Pearson corr; P > 0.05), while coloured ones represent statistically significant correlations (Pearson corr; P < 0.05). False Discovery Rate (FDR) correction was applied for p values using the Benjamini-Hochberg procedure. The size of the circle depicts the degree of significance (the bigger, the smaller the value for P) and the color refers to the direction of the correlation (brown -or left side of the bar- is negatively correlated, green -or right side of the bar- is positively correlated). The icons refer to the nature of the variable (Treatment, plant biomass, leaf, microbiome, or soil-related). MP = Microbiome Predator.
